# Slow Spindle Trains During Daytime Naps are Associated with Improved Declarative Memory Consolidation

**DOI:** 10.64898/2025.12.29.694712

**Authors:** Vaishali Mutreja, Prakriti Gupta, Ovidiu Lungu, Latifa Lazzouni, Arnaud Boutin, Ella Gabitov, Madeleine Sharp, Julie Carrier, Julien Doyon

## Abstract

Memory consolidation refers to the process by which newly encoded memories are strengthened and retained over time, and ample evidence indicates that sleep supports this process for both procedural and declarative memories. Although sleep spindles during non-rapid eye movement (NREM) sleep have been associated to consolidation, it remains unclear whether all spindle types contribute equally. Spindles vary in frequency and topography–slow spindles (≤12.5Hz) predominating over frontal regions, whereas fast spindles (>12.5Hz) peak parietally – and recent work suggests that procedural memory consolidation during overnight sleep is related to the temporal organization of spindles in ‘trains’ (i.e., events occurring <6s apart). Here we investigated whether a similar mechanism operates for declarative memory during daytime naps. Participants were assigned to a Nap (N=23) or No-Nap (N=15) group, and completed an object-spatial location task involving 36 item-location associations. Memory was assessed immediately after learning and again following a 90-minute nap or an equivalent wake period. Results showed that the Nap group exhibited significantly better delayed memory, as measured by combined recall-recognition score, and a greater proportion of participants maintained or improved their performance. In the Nap group, memory performance correlated with local spindle density at frontal and parietal sites, and, critically, with the proportion of slow spindles clustered in trains during NREM2. These findings suggest the temporal organization of slow spindles into clusters support declarative memory consolidation, pointing to a shared spindle-based mechanism across domains.

---

Memory consolidation is the process by which newly acquired memories, initially vulnerable and susceptible to interference, are stabilized and transformed into more durable representations over time (Dudai et al., 2015). A growing body of research has highlighted the crucial role of sleep in facilitating this transformation (Boutin & Doyon, 2020; Gais & Born, 2004; Rasch & Born, 2013). Sleep benefits both declarative memories, which encompass consciously retrievable information such as facts, word-pairs, and spatial association, and procedural memories, which involve the acquisition of different motor skills. Although direct mechanistic evidence in humans remains limited, coordinated neural activity during sleep is thought to promote the reactivation and reorganization of memory traces, enabling their integration into long-term storage (Diekelmann & Born, 2010; Klinzing et al., 2019). Among the various sleep-related oscillatory phenomena, sleep spindles – brief bursts of activity in the sigma frequency range (∼9-16Hz) occurring mainly during non-rapid eye movement (NREM) stage 2 sleep – have emerged as key candidates supporting this process (Boutin & Doyon, 2020; Cox et al., 2017; Diekelmann & Born, 2010; Klinzing et al., 2019). However, a recent meta-analysis reported a stronger and more consistent relationship between spindles and procedural memory (r = .32) than between spindles and declarative memory (r = .21) (Kumral et al., 2023).

Despite substantial progress to date, several important questions remain unresolved in sleep-related memory consolidation research. One key concern is the identification of specific spindle features that contribute to memory consolidation. Rather than all spindles being equally involved in this process, recent work suggests that certain characteristics – such as frequency (slow vs. fast), spatial distribution (frontal vs. parietal), and coupling with slow oscillations – may be more predictive of declarative memory consolidation outcomes (Cairney et al., 2018; Petzka et al., 2022; van Schalkwijk et al., 2019). In particular, slow spindles, which predominate over frontal cortices, have been linked to schema integration, abstraction, and top-down modulation of declarative memory (Helfrich et al., 2018; Matthias Mölle et al., 2011). Yet findings remain inconsistent across studies, possibly reflecting differences in task types (e.g., verbal vs. visuospatial paradigms), analytic approaches (e.g., global vs. local spindle measures), or individual factors such as age, sex and sleep quality, all of which can modulate spindle characteristics and their relation to memory. These sources of variability, however, do not fully account for the discrepancies, suggesting that additional, unexplored spindle features may also play a role.

An emerging line of research based upon work from our own laboratory and others has focused on the temporal organization of spindles into rhythmic clusters known as “trains” – i.e., groups of spindles occurring in close succession less than 6 s apart from each other (Antony et al., 2018; Boutin & Doyon, 2020; Boutin et al., 2024; Solano et al., 2020, 2022). These spindle trains are thought to extend the windows of cortical excitability, potentially allowing for repeated reactivation and stronger consolidation of memory traces (Muehlroth et al., 2019). Although their behavioral relevance has been demonstrated in procedural learning paradigms (Boutin et al., 2024; Solano et al., 2020, 2022), the role of spindle trains in declarative memory consolidation remains largely unexplored.

Also, to our knowledge, no prior study has examined whether the contribution of spindle trains depends on spindle frequency subtypes. Building on prior evidence that slow and fast spindles subserve distinct memory processes – with frontal slow spindles linked to integrative, schema-related operations and parietal fast spindles to hippocampal replay (Helfrich et al., 2018; Lustenberger et al., 2016; Staresina et al., 2015) – it remains unknown whether these frequency-specific mechanisms also differ in their temporal organization. Combining these two dimensions, frequency (slow vs. fast) and temporal clustering (trains vs. isolated), may offer a more comprehensive framework for understanding how spindles coordinate local and distributed neural processes that support memory consolidation. Testing this interaction represents a critical step toward identifying whether distinct spindle subtypes contribute differently when temporally grouped, revealing how spectral and temporal properties jointly shape sleep-dependent memory processes.

A further limitation in the literature concerns the lack of proper control conditions in overnight sleep studies. For the few studies that have included sleep-deprived comparison groups, such control conditions are confounded by factors like fatigue, reduced alertness, and circadian misalignment, making it difficult to isolate the specific contribution of sleep to memory outcomes (Alhola & Polo-Kantola, 2007; Gais et al., 2006; Newbury et al., 2021). In this regard, daytime naps paradigms offer a promising alternative given that a wake control group deprived of a nap is not affected by the above-mentioned confounders. Moreover, naps are shorter in duration, more easily scheduled in controlled settings, and reliably include NREM sleep implicated in declarative memory consolidation (Schabus et al., 2004). Additionally, naps minimize the effects of circadian variability and prolonged wakefulness (Lahl et al., 2008; Tucker & Fishbein, 2008), while still preserving key physiological features of nocturnal sleep, including the presence of sleep spindles that mirror those observed at night (Mylonas et al., 2020). Yet, despite these advantages, naps remain underutilized in studies probing the role of spindles, particularly in the declarative memory domain.

The current study thus sought to address these outstanding issues by investigating whether spindle trains during daytime NREM sleep support declarative memory consolidation of newly learned visuospatial information, measured with an object-location memory task. We quantified the local spindle density across midline derivations (Fz, Cz, Pz) and examined their temporal organization into trains during NREM2 sleep, separately for slow and fast spindle subtypes. While prior studies have examined spindle frequency (slow vs. fast) and temporal organization (train vs. isolated) as separate dimensions, these features may reflect complementary aspects of sleep-dependent consolidation: frequency capturing which neural networks are engaged (frontal vs. parietal), while temporal organization reflecting how often and persistently these networks are reactivated. Combining both dimensions therefore provides a more comprehensive test of whether the temporal clustering of spindles modulates their frequency-specific contributions to memory consolidation. Specifically, we tested whether memory retention was predicted by (1) spindle frequency subtype, (2) temporal organization of spindles, or (3) the interaction of these two dimensions. Based on prior evidence linking spindle trains to repeated memory reactivation and frontal slow spindles to integrative memory functions, our primary hypothesis focused on whether clustering of slow spindle into trains would facilitate the consolidation of newly encoded visuospatial information. Analysis involving fast spindle trains were conducted as secondary, complementary exploratory analyses, aimed at assessing whether temporal clustering effects generalized beyond slow spindles. Clarifying this relationship may enhance our mechanistic understanding of how naps contribute to memory consolidation and inform spindle-targeted interventions in both healthy and clinical populations.

## Methods

### Participants

Fifty-two healthy young adults (20-35 years) were recruited through local advertisements and provided written informed consent. The study was approved by the Research Ethics Board (CMER RNQ 13-14-011) of the ‘Centre de Recherche de I’lnstitut Universitaire de Gériatrie de Montréal (CRIUGM)’ and conducted in accordance with the Declaration of Helsinki (2013) (Shrestha & Dunn, 2020). Participants received monetary compensation. Eligibility criteria required right-handedness, good physical and mental health, absence of sleep disorders, and no use of psychoactive or sleep-affecting medications. Screening included self-report questionnaires assessing handedness, sleep quality, depression, anxiety, and daytime sleepiness; full details are provided in Supplementary Materials. Participants were asked to maintain a regular sleep-wake schedule for at least three days prior to the experiment and to abstain from alcohol, caffeine, and nicotine for 24 hours.

Participants were assigned to either a Nap group (n=31; 21 females; mean age=23.98 ± 3.28 years), who received a 90-minute nap opportunity, or a No-Nap group (n=21; 12 females; mean age=24.76 ± 4.40 years), who remained awake under EEG monitoring. Although participants were aware that the study involved both nap and wake conditions, their individual assignment was revealed only on the experimental day. Fourteen participants were excluded: three for failing to reach the learning criterion, two for having <10 minute of NREM2 sleep, and nine due to missing behavioral data. The final sample included 38 participants, with no significant group differences in age or sex (Table 1), as confirmed by independent samples t-test and chi-square analysis [age (t(36) =1.01, p=.33), sex (c^2^(1) =.13, p=.72)]. Thus, any observed group differences in memory performance are unlikely to reflect demographic factors.

**Table 1.** Participants’ characteristics

| Group | Nap | No-Nap |
| --- | --- | --- |
| Total Participants | 23 | 15 |
| Females | 14 | 10 |
| Males | 9 | 5 |
| Age-range | 20-34 | 20-35 |
| m<sub>age</sub> (SD) | 24.09 (2.84) | 25.47 (4.78) |
| m<sub>age</sub> (SD) Female | 23.50 (2.62) | 24.40 (4.53) |
| m<sub>age</sub> (SD) Male | 25.00 (3.08) | 27.60 (5.03) |

Notes. Values are in years. m<sub>age</sub> = mean age of the participant; SD = Standard deviation

### Overall experimental design and procedure

The study consisted of a between-subjects design with two groups: Nap and No-Nap (Figure 1a). Participants who met the inclusion criteria were invited to our sleep laboratory for the experimental session. After installing the EEG system on all participants (Nap and No-Nap groups), they were trained on a declarative memory paradigm (see below), followed by an immediate cued-recall test and by a 90-minute nap (Nap group) or a wake period for the same amount of time (No-Nap group). After the nap or wake period, participants were required to complete delayed cued-recall and recognition tests.

**Figure 1.**
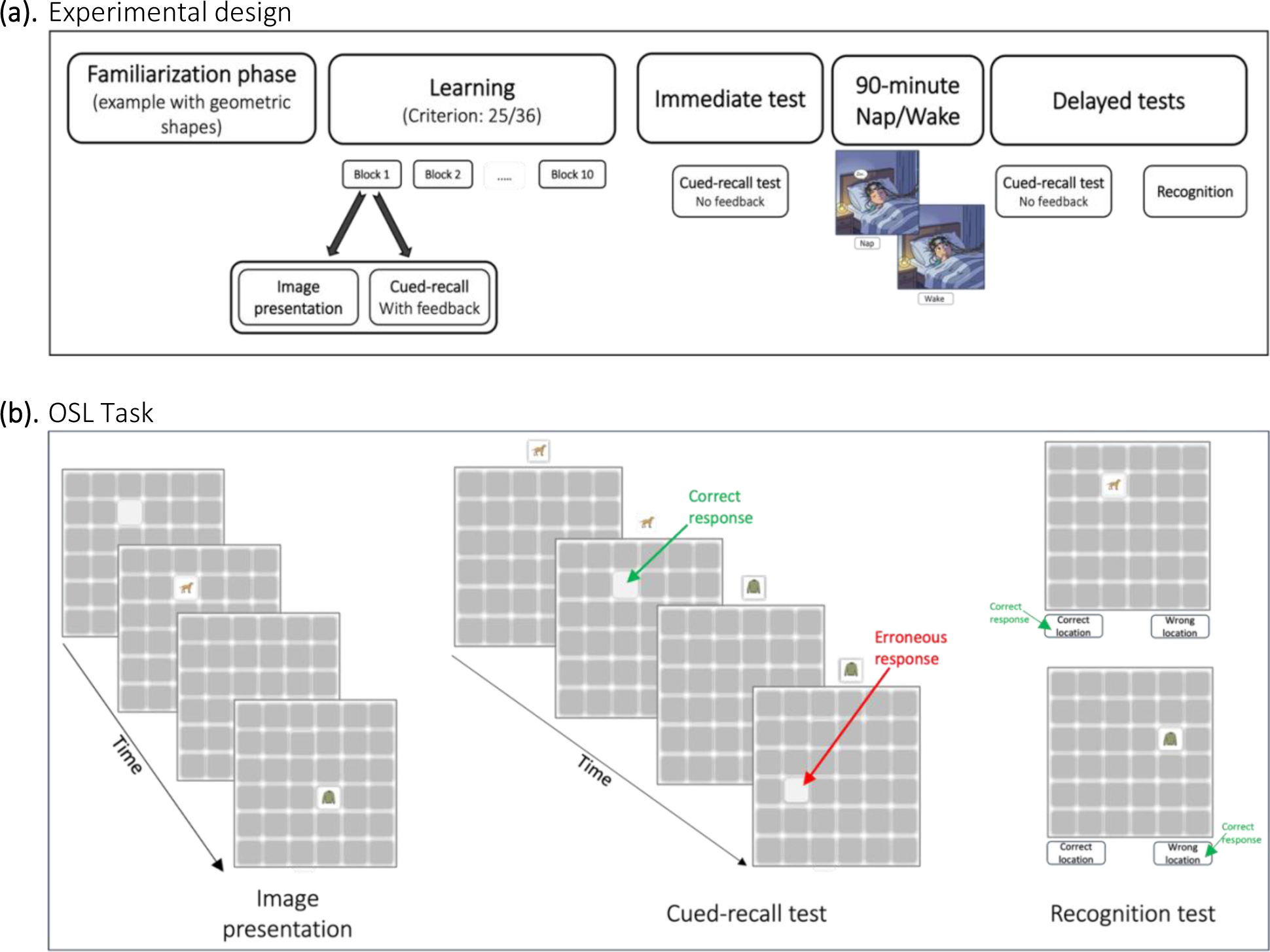
Experimental design and OSL task. (a). *Experimental design*. Participants first completed a short familiarization phase with simple geometric shapes, followed by the learning phase of the OSL task (criterion = 25/36 correct). The learning phase consisted of repeated blocks of image presentation and cued-recall test with feedback. After reaching a learning criterion (70% of correct responses), participants performed an immediate cued-recall test (no feedback) and were then either allowed a 90-minute nap (Nap group) or asked to remain awake (No Nap group); participants in both groups were monitored using polysomnography. Finally, they completed two delayed memory tests (no feedback); a cued-recall test and a recognition test. (b). *OSL Task.* During the learning phase, each block consisted of two parts: an image-presentation phase, where 36 images were shown one at a time in their unique grid locations in a 6×6 grid, and a cued-recall test, where each image appeared above the grid and participants indicated its learned location (feedback provided only during learning). In the recognition test, each image was shown twice–once in the correct location and once in a wrong location–and participants had to indicate whether the location was correct or wrong.

### Object-Spatial Location (OSL) Declarative Memory task

Declarative memory was assessed with a two-dimensional object-spatial location (OSL) task (Figure 1b) adapted from prior visuospatial memory paradigms (Rudoy et al., 2009; Shanahan et al., 2018). Thirty-six colored images (9 per category: animals, clothing, food, vehicles) were each assigned a unique position within a 6×6 grid. Stimuli were selected from the revised Snodgrass and Vanderwart objects set (Rossion & Pourtois, 2004) and randomly placed in a unique position within it. The task was programmed in PsychoPy v3.6 (Peirce et al., 2019).

### Learning and Tests

#### Familiarization phase

Participants first completed a brief familiarization phase to learn the grid structure and response procedure. They practiced associating simple geometric shapes with specific grid locations and advanced after two consecutive blocks without any errors (full procedural details in Supplementary Materials).

#### Learning

Participants then learned the 36 object-location associations through repeated blocks consisting of an image presentation part followed by a cued-recall test with feedback. Blocks continued until participants reached a learning criterion of 70% accuracy or completed a maximum of 10 blocks, whichever occurred first.

#### Immediate memory test

After a short break of 3-minute, participants competed an immediate cued-recall test identical to learning but without feedback.

#### Delayed memory tests

Following a 90-minute nap (Nap group) or equivalent wake period (No-Nap group), participants completed two delayed tests: (a) a delayed cued-recall test – identical to the immediate memory test, and (b) a recognition test assessing whether each image appeared in its correct or an incorrect grid location. Each of the 36 images was shown once in the correct location and once in a different (wrong) location. (Detailed timing and response procedures are provided in Supplementary Materials.)

### Behavioral data analyses

#### Learning phase

During learning, a total score for each block was computed as the number of correctly recalled image locations, to capture participants’ learning curve across the task. In addition, two indices of learning efficiency were calculated: the number of blocks each participant required to successfully complete this phase, and the total score obtained on the last learning block.

#### Cued-recall test

Three variables were used to assess participants’ performance across the immediate and delayed cued-recall tests, hence allowing us to evaluate their memory retention capacity and changes over time: 1) Correct immediate-cued recall score: i.e., the number of image locations accurately recalled during the immediate-cued recall test; 2) Correct delayed cued-recall score: i.e., the number of image location accurately recalled during the delayed cued-recall test; and 3) Retained image locations: i.e., the number of image locations correctly recalled during both immediate and delayed cued-recall tests.

#### Recognition

The participants’ performance during the recognition test was assessed using two variables: 1) Hits: i.e., the number of images correctly identified as being in their correct location in the grid; 2) Correct rejection: i.e., the number of images correctly identified as being presented in a wrong location in the grid. Both hits and correct rejection scores provided insight into the participants’ spatial memory accuracy.

The data derived from learning and recall variables, in combination with those from the recognition test, contributed to calculating the participants’ combined recall-recognition scores (see below).

#### Combined recall-recognition scores

Two dependent composite scores were derived from the above variables: *C1. Combined recall-recognition score* corresponding to the percentage of image locations that were retained in both cued-recall tests, as well as correctly recognized (hit) and correctly rejected image locations presented during the recognition test. This composite score represents an integrated assessment of memory consolidation across both cued-recall and recognition tests. *C2. Memory retention status*: a nominal score capturing whether each participant’s cued-recall performance improved or was maintained vs. deteriorated from immediate to delayed recall tests.

### Polysomnographic recording and analysis

EEG was recorded using a BrainAmp system (Brain Products FmbH) with a 64-channel cap referenced to FCz and grounded at Afz; 22 electrodes positioned according to the 10-20 system were used for analyses. Signals were sampled at 5000 Hz and impedance was kept <5kΩ. Pre-processing was performed in BrainVision Analyzer 2.1: data were down-sampled to 250Hz, re-referenced to M1/M2, band-pass filtered (0.3-30Hz), and notch-filtered at 60Hz. Automatic artifact detection relied on BrainVision default criteria to identify ocular and movement-related artifacts, and flagged segments were marked with boundary events. Sleep stages were scored by a trained polysomnographic technologist using Hume in MATLAB (https://www.jaredsaletin.org/hume) according to the standard criteria (Iber et al., 2007). Spindles were detected automatically on artifact-free NREM2/3 epochs at Fz, Cz, and Pz using a dynamic thresholding algorithm (Boutin & Doyon, 2020; Warby et al., 2014); signals were filtered in the sigma range (≈9-16Hz), and events (0.3-2s) exceeding ∼99^th^ percentile amplitude relative to the RMS of the filtered signal were retained. Additional methodological details are provided in the Supplementary Materials.

### Variables included in EEG analyses

We quantified three spindle characteristics: 1) *spindle density:* global density (spindles/minute of NREM sleep) and local density (spindles within a spindle-centered sliding window of 60s); 2) *proportion of spindles based on their frequency*: slow (≤12.5 Hz) versus fast (>12.5Hz) spindles (Cox et al., 2017); and 3) *proportion of spindles based on their clustering*: isolated spindles (>6s apart) versus spindle trains (<6s apart; Boutin and Doyon (2020)). For each spindle type (i.e., slow and fast), we also computed the proportion occurring in trains. We focused on midline electrodes (Fz, Cz, and Pz) to capture the anterior-posterior distribution of spindle subtypes, with slow spindles being maximal frontally and fast spindles parietally (Andrillon et al., 2015; M. Mölle et al., 2011), and because these sites provide standardized sampling of spindle activity (De Gennaro & Ferrara, 2003).

### Statistical Analyses

All statistical analyses were conducted using the MATLAB-R2021b (MathWorks Co.) and SPSS v29. Before group comparisons, data were assessed for normality (Shapiro-Wilk) and homogeneity of variances (Levene’s test). Parametric tests (independent-samples t-tests) were used when assumptions were met; otherwise, non-parametric tests (Mann-Whitney U) or adjusted tests (e.g., Welch’s t-test) were applied. When both approaches produced convergent results, parametric outcomes are reported.

For behavioral data, we first verified comparable baseline performance by comparing recall accuracy at the end of learning using an independent-samples t-test. Both groups also performed similarly on the immediate recall test prior to the 90-minute nap or wake interval. To test the hypothesis that napping after learning enhances declarative memory consolidation more as compared to staying awake, we used Welch’s t-test (due to unequal variances) to compare the percentage of combined recall-recognition score between the Nap and No-Nap groups. Additionally, we conducted a chi-square test to assessed group differences in the proportion of participants who improved or maintained performance versus those who deteriorated from immediate to delayed recall; participants who maintained their performance (i.e., showed no change) were grouped with those who improved, reflecting the stabilizing role of naps. Finally, to assess whether encoding strength during learning predicted memory retention, Pearson correlations were computed between two learning metrics–the percentage of correct responses in the last block of learning and the number of learning blocks required to reach criterion–and the combined recall-recognition score.

For EEG data (Nap group only), we first computed descriptive analyses of sleep and spindles characteristics during NREM2, including spindle features across frontal (Fz), central (Cz), and posterior (Pz) midline derivations. To examine how spindle activity related to memory consolidation, we conducted Pearson correlations between spindle density measures (global and local) and post-nap memory performance. We then applied a three-tier analytic approach to characterize spindle dynamics: (1) assessing whether overall spindle clustering (proportion of spindles in trains vs. isolated) predicted memory performance; (2) examining correlations between spindle frequency subtypes (slow vs. fast) and memory performance, independent of their temporal organization; and (3) testing the interaction of these dimensions by correlating the proportion of slow and fast spindles occurring in trains with memory performance. Consistent with our a priori hypothesis, the primary analysis focused on slow spindle trains, whereas the fast spindle analyses were treated as complementary exploratory tests. A significant threshold of p<.05 was applied, and one-sided correlations were used given directional predictions of positive associations with memory.

## Results

### Behavioral results

#### Baseline memory performance before a nap or wake period

The independent samples t-test showed no significant difference between the Nap (*M*=80.56, *SEM*=2.77) and No-Nap (*M*=81.65, *SEM*=3.60) groups at the end of the learning, *t*(36) =0.47, *p*=.64; Levene’s test confirmed equal variances (*p*=.83; Figure 2a). Immediate cued-recall performance was also comparable between the Nap (*M*=74.39, *SEM*=4.06) and No-Nap (*M*=77.04, *SEM*=4.49) groups, *t*(36) =0.85, *p*=.39, with equal variances (*p*=0.64). A secondary Welch’s t-test confirmed this similarity *t*(33.248) =0.88, *p* =.39. Thus, both groups exhibited equivalent baseline memory performance prior to the 90-minute nap or wake period.

**Figure 2.**
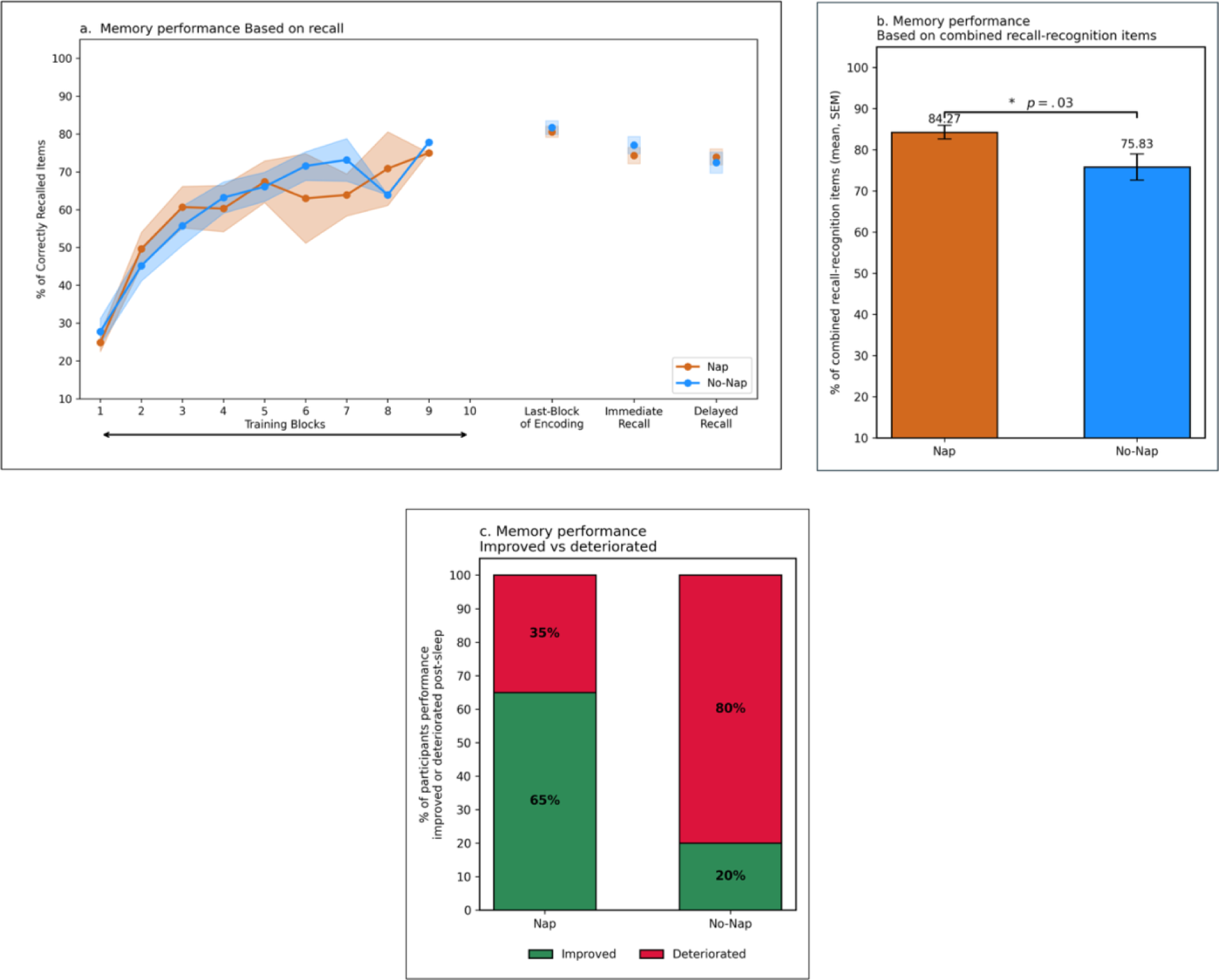
Behavioral results. a) Learning curves: Mean percentage of correctly recalled image locations across learning blocks (1-10), with performance at last block, immediate cued-recall test (10 min post-learning), and delayed cued-recall test (after a 90-min nap or wake interval). b) Consolidation outcome: Mean combined recall-recognition score (retained, correctly recognized, and correctly rejected) in the Nap vs. No-Nap group. * Indicates p < 0.05. Error bars represent standard error of mean (SEM). c) Memory change classification: Proportion of participants in each group who improved or maintained their memory performance (green) vs. those who deteriorated (red) from immediate to delayed cued-recall test.

#### Role of nap in memory consolidation

The Welch’s t-test revealed that the Nap group exhibited a significantly higher combined recall-recognition score (*M*=84.27, *SEM*=1.45) compared to the No-Nap group (*M*=75.83, *SEM*=1.87), Welch’s *t*(21.52) =2.35, *p* =.03 (see Figure 2b). This finding was also corroborated by the chi-square test which revealed that a significantly higher proportion of participants in the Nap group (65%) improved or maintained their performance from immediate to delayed recall compared to the No-Nap group (20%), χ^2^(1, *N*=38) =7.45, *p* <0.001 (Figure 2c), again indicating that post-learning nap increases the likelihood of memory preservation and improvement relative to an equivalent wake period.

### EEG results

Sleep architecture: Participants in the Nap group slept on average of 68.09 minutes (*SEM*=2.70), including 35.96 minutes (*SEM*=2.32) of NREM2 sleep (Table 2a).

**Table 2.** Sleep architecture and sleep spindle characteristics. Sleep architecture and sleep spindle characteristics (mean and SEM) for spindles at the following scalp derivations Fz, Cz, and Pz during NREM2 sleep stage

|  | Mean | SEM | MIN | MAX |
| --- | --- | --- | --- | --- |
| **a Sleep stages** |  |  |  |  |
| Total sleep time | 68.09 | 2.70 | 31.00 | 84.50 |
| NREM1 | 9.20 | 1.22 | 1.0 | 22.5 |
| NREM2 | 35.96 | 2.32 | 16.5 | 56.5 |
| NREM3 | 22.93 | 3.24 | 0 | 47.0 |
| NREM2 + NREM3 | 58.89 | 3.03 | 20 | 81.0 |
| **b Spindles recorded on Fz** |  |  |  |  |
| Number | 192.43 | 17.92 | 63 | 350 |
| % Spindles in trains | 80.31 | 2.23 | 58.18 | 94.57 |
| % Slow spindles | 41.49 | 1.73 | 26.80 | 57.14 |
| % Slow spindles in trains | 54.72 | 2.86 | 30.77 | 74.15 |
| % Fast spindles | 58.51 | 1.73 | 42.86 | 73.20 |
| % Fast spindles in trains | 63.33 | 2.98 | 37.04 | 89.87 |
| Global spindle density | 5.61 | 0.57 | 2.00 | 12.90 |
| Local spindle density | 9.34 | 0.72 | 4.69 | 17.95 |
| **c Spindles recorded on Cz** |  |  |  |  |
| Number | 212.00 | 22.09 | 70 | 401 |
| % Spindles in trains | 80.67 | 2.70 | 38.89 | 98.25 |
| % Slow spindles | 31.32 | 1.79 | 12.50 | 45.21 |
| % Slow spindles in trains | 51.03 | 3.63 | .00 | 87.50 |
| % Fast spindles | 68.68 | 1.79 | 54.79 | 87.50 |
| % Fast spindles in trains | 70.44 | 2.94 | 36.51 | 95.44 |
| Global spindle density | 6.30 | 0.82 | 2.32 | 20.05 |
| Local spindle density | 10.09 | 1.06 | 3.50 | 27.93 |
| **d Spindles recorded on Pz** |  |  |  |  |
| Number | 203.83 | 22.56 | 44 | 379 |
| % Spindles in trains | 80.07 | 3.16 | 27.27 | 98.41 |
| % Slow spindles | 24.77 | 2.37 | 4.55 | 49.15 |
| % Slow spindles in trains | 42.50 | 3.95 | .00 | 85.81 |
| % Fast spindles | 75.23 | 2.37 | 50.85 | 95.45 |
| % Fast spindles in trains | 72.64 | 3.59 | 23.81 | 96.96 |
| Global spindle density | 6.05 | 0.83 | 1.42 | 18.90 |
| Local spindle density | 9.58 | 0.94 | 2.41 | 23.37 |

Notes. Sleep stage values are in minutes. Abbreviations: SEM = Standard Error of Means; MIN = Minimum sleep duration; MAX = Maximum sleep duration, NREM = Non-Rapid Eye Movement sleep, NREM1 = sleep stage 1, NREM2 = sleep stage 2, NREM3 = sleep stage 3.

Sleep spindle characteristics: Spindle counts, density measures, slow/fast spindle distributions, and the proportion of spindles occurring in trains at Fz, Cz, and Pz are summarized in Table 2b–d and reflect expected regional variability in spindle features.

### Role of local spindle density and memory consolidation

Local spindle density correlated positively with combined recall-recognition score at Pz, *r*(21) =0.39, *p*=.03, and at Fz, *r*(21) =0.36, *p*=.05 but not at Cz. Global spindle density at any site was unrelated to combined recall-recognition score (Figure 3). These findings suggest that local spindle density in specific brain regions (frontal and posterior) during NREM2 may be more closely associated with memory consolidation than global spindle density, hence supporting the hypothesis that region-specific spindles contribute to memory consolidation.

**Figure 3.**
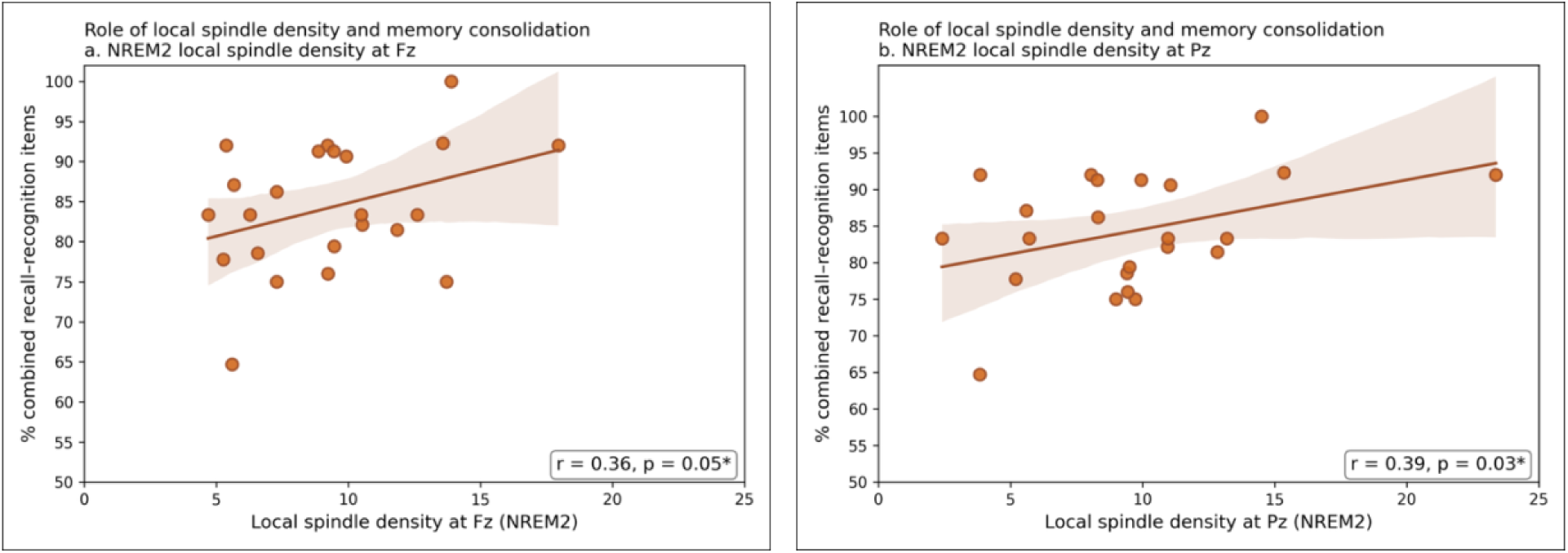
Role of local spindle density and memory consolidation. Correlation between local spindle density (mean number of spindles within a spindle-centered sliding window of 60 seconds) and memory performance (based on combined recall-recognition score) at the frontal (Fz) (left panel) and parietal (Pz) (right panel) midline derivation during NREM 2 sleep. Each panel shows individual data points, linear regression lines, and 95% confidence intervals. Pearson correlation coefficient (*r*) and significance level (*p*-values) are reported for each correlation. NREM: Non-Rapid Eye Movement.

### Spindle clustering in trains and memory consolidation

Across electrodes (Pz, Cz, Fz), the overall proportions of spindles in trains did not correlate with combined recall-recognition score (all *p*s >.07). Similarly, spindle subtypes considered independently of temporal clustering (slow vs. fast) were not associated with combined recall-recognition score (all *p*s >.20). However, when combining both spindle features, a more specific pattern emerged: at Fz, the proportion of slow spindles occurring in trains correlated positively with combined recall-recognition score, *r*(21) =0.35, *p*=.05, (Figure 4, left), whereas fast spindles grouped in trains were unrelated to the combined recall-recognition score, *r*(21) =0.19, *p* =.20 (see Figure 4, right panel). No significant associations were found at Cz or Pz for either spindle subtypes with combined recall-recognition score (all *p*s >.20).

**Figure 4.**
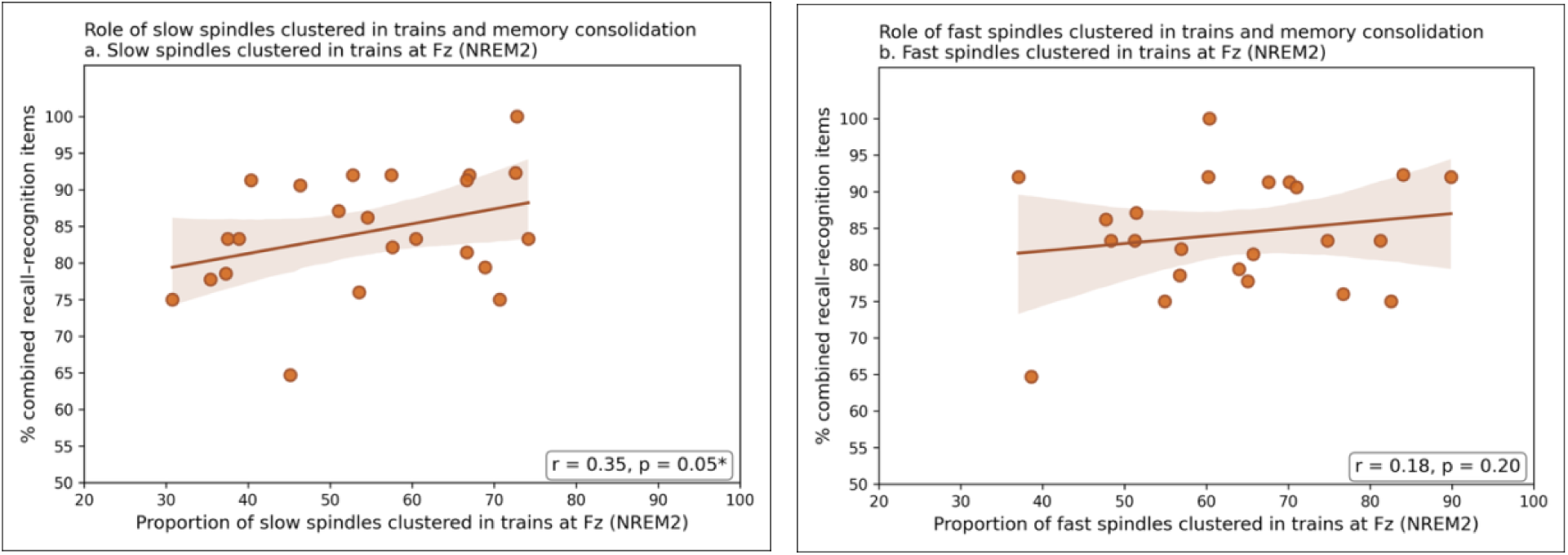
Role of slow/fast spindles clustered in trains and memory consolidation. Correlation between the proportion of slow spindle clustered in trains (left panel) and fast spindles clustered in trains (right panel) at the frontal (Fz) midline derivation during NREM2 sleep and memory performance (percentage of combined recall-recognition items). Each panel shows individual data points, linear regression lines, and 95% confidence intervals. Pearson correlation coefficient (*r*) and significance level (*p*-values) are reported for each correlation. NREM: Non-Rapid Eye Movement.

## Discussion

The present study examined whether a daytime post-learning nap, relative to an equivalent wakefulness period, facilitates the consolidation of visuospatial declarative memory, and whether specific sleep spindle characteristics captured via EEG contribute to this benefit. Behaviorally, the Nap group demonstrated significantly greater memory retention than the No-Nap group, despite comparable performance at baseline, indicating that the advantage attributable to the difference in experimental conditions in the post-learning sleep/wake interval rather than in encoding disparities. Electrophysiologic ally, a selective pattern emerged: only slow spindles clustered in trains at the frontal site (Fz) were positively associated with memory performance. In contrast, neither overall spindle trains nor global proportions of slow spindles–when considered independent of temporal clustering–predicting memory performance. Together, these findings suggest that the efficacy of sleep-based memory consolidation is not uniformly explained by spindle activity but instead depends on the spectral and temporal properties of spindles as well as their regional expression, highlighting that not all spindles contribute equally to memory consolidation.

Our behavioral results are consistent with prior work showing that even brief periods of sleep can benefit declarative memory consolidation (Born & Wilhelm, 2012; Payne et al., 2015). For example, Tucker and Fishbein (2008) reported that a 90-minute nap enhanced performance on a spatial task involving maze navigation, while Wamsley et al. (2010) reported that a nap following virtual maze learning led to significantly greater performance gains than an equivalent wake period. These findings suggest that NREM-rich naps can support the stabilization of hippocampal-dependent spatial traces. In our study, both groups reached comparable levels of learning during the learning phase. Moreover, the individual differences in encoding strength – as measured by the number of correctly recalled image locations during the last learning block and the number of learning blocks needed to achieve the learning criterion (70% accuracy) – did not correlate with memory retention scores. This pattern indicates that the observed nap-related memory benefit is unlikely to be driven by initial encoding differences. The significant proportion of participants who showed memory gains or performance maintenance in the Nap group, together with converging evidence from previous nap studies showing similar enhancements in declarative memory (e.g., Lahl et al. (2008); Tucker and Fishbein (2008); Wamsley et al. (2010)) are consistent with an active memory consolidation mechanism – in which sleep promotes the stabilization and long-term retention of memory – exceeding what can be explained by passive protection from waking interference alone.

EEG analyses revealed that the nap-related memory benefit was associated with region-specific spindle dynamics. In line with the view that spindles are locally rather than globally synchronous events (Tononi & Cirelli, 2006), our results showed that local spindle density at both frontal (Fz) and posterior (Pz) sites was positively associated with the visuospatial memory retention. This finding echoes prior work by Clemens et al. (2006), who found that visuospatial memory consolidation was selectively associated with spindle counts over parietal regions, whereas verbal memory benefits were linked to spindles over left frontocentral areas. Such topographical specificity suggests that spindle activity may reflect the reactivation of task-relevant cortical networks involved during initial encoding.

While our findings focused on cortical signatures captured through EEG, spindles are known to be initiated in subcortical thalamic circuits and relayed to the cortex via thalamocortical loops (Steriade et al., 1993). Thus, the regional effects we observe at the scalp likely reflect coordinated interactions between deep and surface-level generators. Multimodal approaches, such as simultaneous EEG-fMRI or intracranial recordings, may help elucidate how scalp-level spindle activity reflects deeper subcortical-cortical interactions involved in memory consolidation (e.g., (Baena et al., 2024; Chen et al., 2025; Fang et al., 2017).

Our results with regional local density highlight the importance of temporal organization of spindles – specifically their occurrence in rhythmic clusters or “trains”. We observed that only slow spindles occurring in clusters at frontal sites were significantly associated with enhanced memory performance. This finding complements prior work by (Boutin et al., 2024) who showed that spindle trains during NREM2 sleep predicted motor memory gains, suggesting that spindles trains may provide repeated opportunities for memory reactivation, beyond what the isolated spindle events can offer. Theoretical models propose that such clustering prolongs periods of heightened cortical excitability and plasticity (Antony et al., 2019), creating multiple windows for memory trace reactivation. In line with this, our findings extend the relevance of spindle trains to the declarative domain and suggest that their presence may reflect an underlying neural architecture optimized for systems-level consolidation. Although the mechanisms driving this temporal organization remain under investigation, accumulating evidence supports the view that spindle trains may scaffold memory reactivation by pacing neural replay across successive cycles (Antony et al., 2018; Antony et al., 2019; Boutin & Doyon, 2020; Fernandez & Lüthi, 2020).

Building on the findings of Boutin et al. (2024) – who demonstrated that spindle trains predicted procedural learning gains–our results extend this framework to declarative memory and reveal a frequency-specific refinement. The association emerged exclusively for slow spindles (∼9-12.5 Hz) organized into trains at the frontal site (Fz), but not for fast spindle trains (>12.5-16 Hz) at parietal sites (Pz), suggesting that the temporal benefit of clustering depends on the neural circuit engaged. This pattern likely reflects the unique architecture of fronto-hippocampal networks, where slow frontal spindles often coupled with slow-oscillation upstate, promote schema integration and top-down transformation of memories (Helfrich et al., 2018; M. Mölle et al., 2011). By contrast, fast parietal spindles are more directly tied to hippocampal replay and item-specific reinstatement (Lustenberger et al., 2016; Staresina et al., 2015) – mechanisms more critical for procedural learning. The NREM2-dominant composition of naps may further favor this frontal slow-spindle expression, providing an optimal milieu for integrative rather than replay-based consolidation. Moreover, clustering slow spindles into trains likely extends successive windows of cortical excitability, creating repeated opportunities for this hippocampal-prefrontal dialogue and gradual stabilization of visuospatial representations (Antony et al., 2018). Thus, while prior work established the general importance of spindle trains, the present findings refine this view, showing that in declarative memory it is specifically slow, frontal spindles trains that orchestrate consolidation – linking frequency, timing, and memory function within a unified framework.

Our finding that only slow spindles clustered in trains predict behavioral memory gains further supports that the memory-related impact of spindles depends on *both* their temporal dynamics and frequency-specific characteristics. While fast spindles may support the initial hippocampal reactivation of encoded traces, however it is slow spindle trains over frontal regions that appear to provide the temporal scaffolding necessary for top-down integration and long-term transformation of memory representations across distributed cortical networks. This spectral and temporal specificity reinforce the emerging view that sleep-dependent consolidation is depends jointly on when and where spindles occur, and moreover how they unfold over time.

### Limitations and Future directions

While the present study provides important insights into how naps and spindle dynamics support the consolidation of visuospatial declarative memory, several limitations should be noted. First, we did not include pre-experimental sleep -wake monitoring or a habituation nap, which would have enabled more systematic screening of sleep difficulties. Although no issues emerged during the experiment, this remains a methodological constraint. In addition, because many participants obtained very little NREM3 sleep, we were unable to examine slow-oscillation-spindle coupling – an important mechanism implicated in overnight systems-level consolidation (Helfrich et al., 2018; M. Mölle et al., 2011; Staresina et al., 2015). Further studies using longer nap opportunities, full-night recordings or prior sleep-wake monitoring may clarify how pre-sleep conditions and slow oscillations–spindles interactions contribute to declarative memory stabilization.

Second, although our EEG was recorded from 22 electrodes, spindle analyses focused on midline sites (Fz, Cz, Pz), chosen a priori to capture frontal slow and parietal fast spindle activity. While source reconstruction would have been feasible, the reduced electrode density would have limited spatial precision. High-density EEG or multimodal approaches (e.g., EEG-fMRI) could extend these findings by identifying spindle generators with greater anatomical specificity and examining their interactions with broader hippocampal-cortical networks.

Finally, the associations between spindle characteristics and memory performance are correlational in nature, hence limiting causal interpretations. To address this, future studies could employ closed-loop targeted memory reactivation–timing cues to specific spindle subtypes or train configurations—may allow more direct testing of the causal role of spindle trains in memory consolidation and could inform development of spindle-targeted interventions in populations with memory impairments.

## Conclusion

In conclusion, this study demonstrates that a brief daytime nap supports the consolidation of visuospatial declarative memory and that this benefit is specifically linked to slow spindles grouped into trains. These findings underscore the importance of both the topographical and temporal architecture of sleep oscillations in understanding how sleep supports memory consolidation. Notably, the results point to a potentially unique role of slow frontal spindle trains in facilitating systems-level integration processes. Taken together, our findings contribute to a more nuanced view of sleep’s role in memory–one of which the *where*, *when* and *how* of sleep oscillations jointly determine the success of consolidation.

## Author contributions

*Vaishali Mutreja*: Writing – original draft, conceptualization, data collection, methodology, formal analysis, figure generations, data curation; *Prakriti Gupta*: Data collection, writing – review and editing; *Ovidiu Lungu*: Conceptualization, methodology, data collection, data analysis, writing – review and editing; *Latifa Lazzouni*: Conceptualization, methodology, data collection, data analysis, writing – review and editing, data curation; Arnaud Boutin: Writing – review and editing; *Ella Gabitov*: Conceptualization, methodology, review and editing; *Madeleine Sharp*: Conceptualization, Writing – review and editing; *Julie Carrier*: Conceptualization, Writing – review and editing; *Julien Doyon*: Funding acquisition, conceptualization, methodology, writing – review and editing, supervision.

## Acknowledgements

This work was supported by the Canadian Institutes of Health Research (CIHR). We thank all participants for their time and commitment to the study. We are also grateful to Hannah James for assistance with data collection. Finally, we thank our colleagues in the SPINDL Laboratory for their helpful discussions and support throughout the development of this project.

## Data Availability Statement

The data that support the findings of this study are available from the corresponding author upon reasonable request

