## Supplemental file for "Slow Spindle Trains During Daytime Naps are Associated with Improved Declarative Memory Consolidation"

#### Supplementary Material

### Methods

#### Participants

Healthy young volunteers (20-35 years) were recruited through local advertisements. Screening was conducted through an online LimeSurvey questionnaire followed by in-lab verification. To be eligible for the study, participants were required to be right-handed (Edinburgh Handedness Inventory; Oldfield (1971)), b) non-smoker, c) non-drug users, d) free of medical, neurological, psychological or psychiatric conditions, including depression and anxiety (score ≤8 on the Beck Depression (Beck et al., 1974) and Anxiety inventories (Beck et al., 1988)), e) free of sleep disturbances (assessed using the Pittsburgh Sleep Quality Index Global Score<5 (Buysse et al., 1989)), and f) free of psychoactive or sleep-affecting medications as per self-reporting. Criteria b, c, and d were assessed through questions included in an online LimeSurvey questionnaire.

Participants were excluded if they self-reported being shift workers, having had experienced a trans-meridian trip less than three months prior to the experiment, or exhibiting signs of excessive daytime sleepiness (≥ 9 on the Epworth Sleepiness Scale (Johns, 1991)). Participants were asked to maintain a regular sleep-wake cycle (bedtime between 10:00-11:30pm, wake-time between 07:00-08:30am) for at least three days before the experiment. They were also required to abstain from consuming alcohol, caffeine, or nicotine at least 24 hours prior to the experiment.

A total of 52 participants were enrolled and assigned to either the Nap (n = 31) or No-Nap (n = 21) group. Assignment to the nap or wake condition was disclosed only on the experimental day, although participants were informed during recruitment that both conditions existed. In the Nap group, participants received a 90-minute nap opportunity, whereas participants in the No-Nap group remained awake in a lighted room under EEG monitoring to ensure wakefulness. Fourteen participants were excluded from the analysis: three for failing to reach the learning criteria, two for having <10 minutes of

NREM2 sleep, and nine due to missing behavioral data caused by software issues. This resulted in a final sample of 38 participants used for behavioral and EEG analyses. Demographic comparisons between groups (reported in the main text) confirmed that the Nap and No-Nap group did not differ significantly in age or sex.

#### Object-Spatial Location (OSL) Declarative Memory task: Detailed Procedures

Declarative memory was assessed using a two-dimensional object-spatial location (OSL) task programmed in PsychoPy v3.6 ((Peirce et al., 2019). The task included 36 colored images drawn from four semantic categories (animals, clothing, food and vehicles; 9 images per category). All stimuli were selected from the revised Snodgrass and Vanderwart image set (Rossion & Pourtois, 2004) and were each assigned a unique position within 6x6 grid. Image locations were randomized across participants.

**Familiarization phase**: The experiment began with a familiarization phase, the purpose of which was twofold: (a) to familiarize participants with the grid layout and (b) to ensure that they understood the task of associating an image with a specific location prior to engaging in the main experiment, which involved learning 36 images and associating each of them with a unique grid location. Participants were presented with three geometric shapes (a disc, a square, and a triangle), one at a time, in the 6x6 grid. Each image was displayed for 2000ms in a specific location. Participants were instructed to memorize the location of the shapes as they were sequentially displayed. After the shapes were all presented, one of the three shapes reappeared at the top of the grid, again for 2000ms. The image then disappeared, and participants had a maximum of 5000ms to indicate (by clicking with a computer mouse) its corresponding location in the grid. This process was repeated for all three shapes until participants completed correctly two consecutive blocks without any errors. After each block, participants received feedback indicating whether all shapes had been correctly located or not.

**Learning**: Following the familiarization phase, participants started the learning phase, which consisted of blocks, each containing two parts: (a) images presentation part and (b) cued-recall test. During the image presentation part, the 36 images from the four categories (animals, clothing, food, and vehicles) were presented one at a time for 2000ms each, in their unique location in the grid. Participants were instructed to memorize the location of each image. Once all images were presented, the cued-recall test began. It consisted of 36 trials – one for each image. On each trial, one of the images appeared on top of the grid for 2000ms and, similar to the familiarization phase, participants had a maximum of 5000ms after the image disappeared to indicate its location in the grid as they remembered it. Participants received feedback at the end of the cued-recall test in the form of a numerical score indicating the number of images they located correctly (e.g., '18/36'). After completing each learning block (the image presentation part and its subsequent cued-recall test), participants were asked to rest for 25s. The learning blocks were repeated until participants either reached a learning criterion of 70% accuracy (i.e., '25/36') or completed a maximum of 10 learning blocks, whichever occurred first.

**Immediate memory test**: Participants who successfully completed the learning phase were given a 3-minute break before starting the immediate cued-recall memory test. This test was identical to the cued-recall tests during the learning phase, except that this time no feedback was provided.

**Delayed memory tests**: Following either a 90-minute nap (Nap group) or wake period (No-Nap group), each participant each completed two tests: (a) a delayed cued-recall test – identical to the immediate cued-recall test, and (b) a recognition test. No feedback was provided during these tests.

The recognition test served two purposes: (a) to assess the participants' ability to recognize previously learned image location pairs, as a complement to the more effortful retrieval process required during cued recall, and (b) to capture potential memory traces that may not have been accessible via recall, thereby offering a complementary index of memory retention. During this test, participants were presented with the image in one of the locations in the grid and were asked to indicate whether it appeared in the correct location or wrong location as per the learning phase. During this test, each of the 36 images was presented twice, once in the correct location and once in a different (wrong) location. Each image was presented for 2000ms and after it disappeared, participants had up to 5000ms to respond by choosing one of two possible options displayed at the bottom of the display screen: 'Correct location' and 'Wrong location'. Participants were instructed to select 'Correct location' if they believed that the image appeared in the correct location (as presented during the learning phase) or 'Wrong location' if they believed it appeared in a wrong location.

#### Polysomnographic recording and analysis

**EEG data acquisition.** EEG was collected using a 64-channel BrainAmp/BrainCap MR system (Brain Products GmbH, Germany), including an ECG electrode. Electrodes (5-kΩ safety resistors) were referenced to FCz with AFz as ground. Twenty-two scalp electrodes from the 10-20 montage were selected for sleep scoring and spindle analyses. Data were sampled at 5000 Hz (500-nV resolution), with impedance maintained below 5 kΩ. Real-time visual inspection ensured stable signal quality throughout the session.

**Pre-processing.** Pre-processing was performed in BrainVision Analyzer 2.1. Signals were down-sampled to 250 Hz, re-referenced to M1/M2, and filtered (0.3-30 Hz) with a 60-Hz notch. Automatic artifact detection relied on BrainVision's default settings: (a) ±200 μV min-max amplitude threshold at Fp1.Fp2 for vertical eye blinks and F7/F8 for lateral eye movements, (b) 50 μV gradient threshold for movement-related artifacts. Flagged segments (ocular activity, movement artifacts, abrupt voltage changes) were marked with "boundary" events to preserve temporal continuity (Delorme & Makeig, 2004).

**Sleep scoring.** A registered polysomnographic technologist scored all recordings using the Hume MATLAB toolbox, following AASM guidelines (Iber et al., 2007). Epochs were classified into NREM1-3, REM, or wake.

**Spindle detection algorithm.** Spindles were detected on artifact-free 30-s NREM2/3 epochs using a dynamic thresholding procedure (Boutin & Doyon, 2020; Gais & Born, 2004; Warby et al., 2014). Signals were band-pass filtered in the sigma band (~9-16 Hz). Spindle candidates (0.3-2 s duration) were defined as oscillatory events exceeding an amplitude threshold set at 0 SD above RMS, equivalent to ~99$^{th}$ percentile – a conservative criterion designed to prevent false detections from background sigma fluctuations or noise.
